# A simple analytical model for predicting locomotive ground reaction forces in foals

**DOI:** 10.1101/2024.12.09.627605

**Authors:** Melany D. Opolz, Sara G. Moshage, Annette M. McCoy, Mariana E. Kersh

**Affiliations:** Department of Chemical and Biomolecular Engineering, University of Illinois Urbana-Champaign; Department of Mechanical Science and Engineering The Grainger College of Engineering, University of Illinois Urbana-Champaign; Department of Veterinary Clinical Medicine, University of Illinois Urbana-Champaign; Beckman Institute for Advanced Science and Technology, University of Illinois Urbana-Champaign; Carle Illinois College of Medicine, University of Illinois Urbana-Champaign

**Keywords:** Froude number, ground reaction forces, equine biomechanics, analytical model, growth

## Abstract

Equine models are useful in biomechanics research of locomotion due to their similarity in musculoskeletal tissue to humans, their athletic nature, and rapid skeletal development which permits ontogenetic studies. However, a continuing challenge in musculoskeletal models for large animal biomechanics is measuring the ground reaction force (GRF) during locomotion. This gap has resulted in a lack of reporting of gait measures such as joint torques. Here we propose an analytical method for predicting ground reaction forces in foals (growing horses) based on the Froude number. Motion capture, GRF, and subject mass data during walking and trotting gaits were collected longitudinally. To account for differences in subject size, we calculated the dimensionless Froude number (Fr=v^2^/(g*l)). The walk-trot transition speed occurred near Fr=0.5, v=1.75-2.15 m/s and was consistent for all evaluated ages. Of the analytical regression models tested, linear regression models had the best performance for predicting vertical GRF data in foals with an average absolute error percent of 7.97% during trotting and 2.39% during walking, compared to a non-linear logistic model with an error of 8.38%. The model was converted to Python and implemented to predict GRF data for foals using the subject velocity and limb length as input. Our resulting analytical model can be used to estimate the GRF profile of equine gait enabling comparative studies of locomotion.

## 1. Introduction

Interest in equine musculoskeletal biomechanics is motivated in part by the similarity in musculoskeletal ailments that affect both horses and humans. The athletic nature of horses [Bryan et al., 2017] leads to orthopedic injuries with similar tissue level manifestations as those found in human injuries [McIlwraith et al., 2011; Rogers et al., 2021]. Thus, the horse serves as a dual-benefit model system [McIlwraith et al., 2012; Kajabi et al., 2021] and has been used in studies of the etiology and treatment of as osteoarthritis [McIlwraith et al., 2012; Kajabi et al., 2021], tendinitis [Henninger et al., 1992; Schramme et al., 2010; Gaesser et al., 2021], and bone stress injuries [Verheyen et al., 2006; Tidswell et al., 2008]. Studies of ontogenetic development are also of interest in the biological, anthropological, and medical sciences. For example, a better understanding of the causes of injuries that occur during growth (stress fractures, growth plate injuries) would allow for improved diagnosis and interventions, but within-subject longitudinal studies of human musculoskeletal development remain impractical. The horse is an ideal model system to circumvent the challenges in ontogenetic studies because they reach skeletal maturity within 2-3 years of age [Rogers et al., 2021].

For many of the musculoskeletal research questions of interest, key outcome measures include: (1) whole body kinematics, (2) musculoskeletal dynamics via joint torques, and (3) the subsequent loading of tissues such as muscle, tendon, and bone. Biomechanical measurements have been enabled by the development of both experimental and computational biomechanics methods, and has resulted in a significant increase in equine musculoskeletal research with studies of equine gait increasing six-fold in the last twenty years. Central to the characterization of joint dynamics and tissue loading during locomotion is the collection of both motion and ground reaction force data.

Ground reaction forces (GRFs) are crucial for characterizing movement and are used to calculate joint torques [Winter, 2009; Koopman et al., 1995; Liu et al., 2009], describe muscle activation patterns [Wakeling et al., 2001; Turns et al., 2007], propulsion and braking during physical activity [Ciacci et al., 2010; Makino et al., 2023], and serve as a quantitative indicator of gait pathologies [Dow et al., 1991; Clayton et al., 2000; Fanchon and Grandjean, 2007; Muniz and Nadal, 2009]. Options for measuring GRFs in equine locomotion include instrumented shoes [Roland et al., 2005], instrumented treadmills [Weishaupt et al., 2002], pressure sensitive insoles [Perino et al., 2007], and force plates [Merkens et al., 1986]. While it is clear that technologies exist for measuring equine GRFs, their use is minimal and is partly attributable to a lack of access to specialized equipment. Consequently, assessments of musculoskeletal functional gait measures, such as joint torques, are sparse with only 13 publications returned in PubMed using the keywords: “equine joint torque” [U.S. National Library of Medicine, nd].

The benefits of equine research for both human and animal health could be better realized by reducing the experimental burden required to collect biomechanics data. Motion capture technologies continue to emerge and advances in inertial measurement units, coupled with machine learning methods, may eventually provide solutions for measuring ground reaction forces. Because of the intrinsic physical relationship between segment kinematics and whole body kinetics (i.e. GRFs), we hypothesize that an analytical model for predicting the ground reaction force that accounts for subject size and speed during steady-state gait may be possible. Such a model would be useful as a first order estimate of the ground reaction force profile for use in comparative studies of musculoskeletal equine biomechanics. Therefore, the aims of this study were to (1) analyze the GRFs in juvenile horses ranging from 2 to 14 months of age in relation to their size, gait type, and speed, and (2) develop and evaluate an analytical model for predicting GRF data in foals.

## 2. Materials and Methods

### 2.1. Subjects

Following IACUC approval from the University of Illinois Urbana-Champaign (UIUC), three male Standardbred trotter foals free of lameness were used for this study. All foals were born at the UIUC Horse Farm and raised on pasture. Longitudinal data collection was performed from 2 to 14 months of age (Fig. 1A). An average of 44 trials were collected per foal per session.

**Figure 1.**
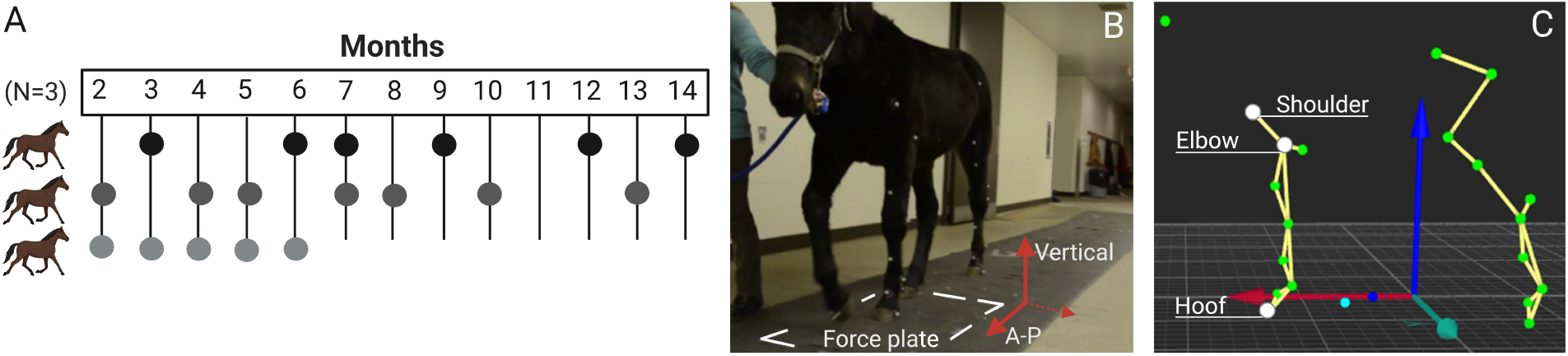
(A) Longitudinal data collection timeline for 3 foals including ground reaction force, motion capture, and mass recordings in each point. (B) One walking trial performed in the hallway inside the animal hospital at UIUC. (C) Marker positions during gait. Markers labeled (white) were used to calculate limb length and velocity.

**Figure 1.**
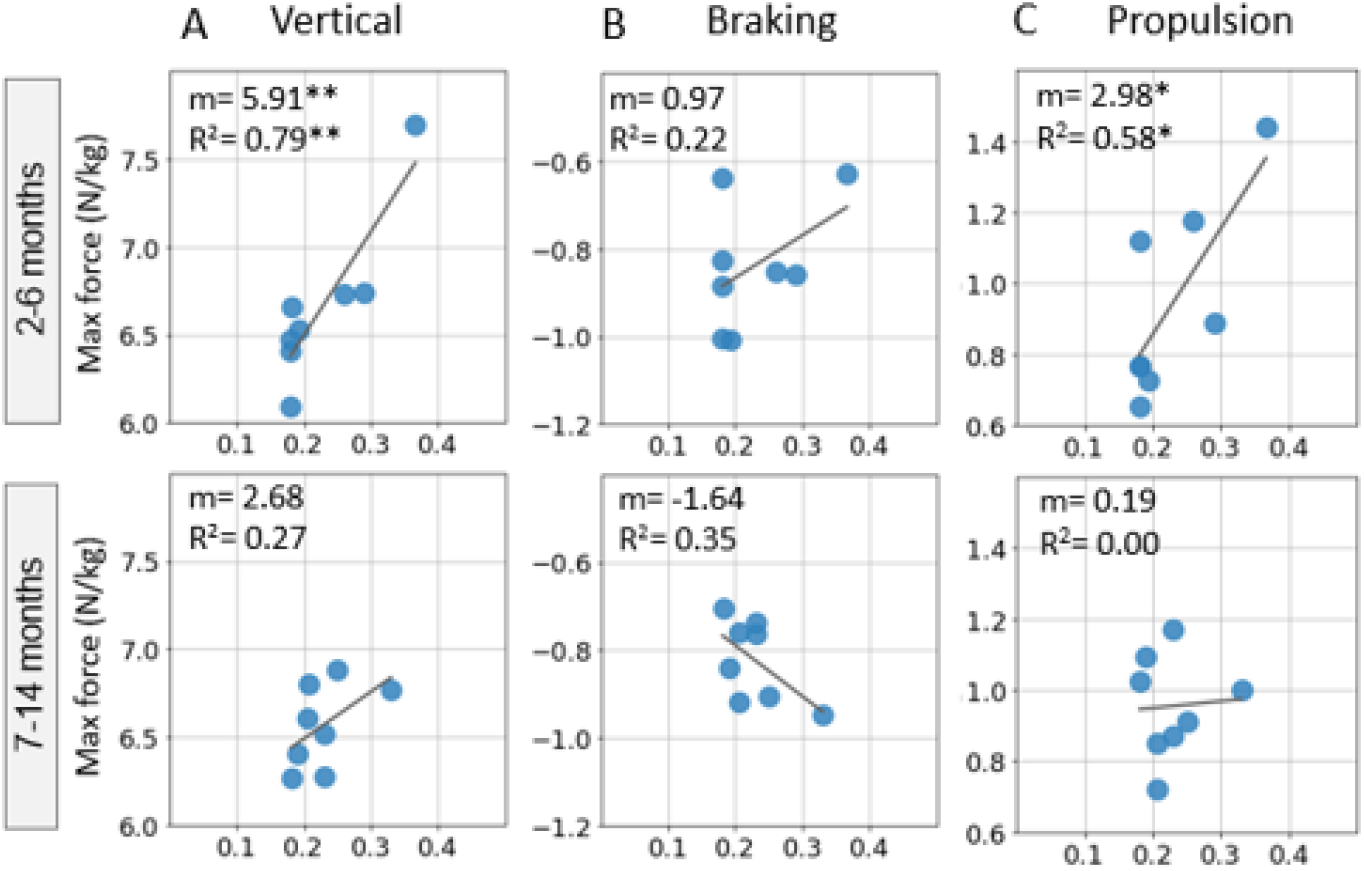
Linear regressions for maximum vertical, maximum propulsion, and maximum braking forces related to Froude number grouped by months according to developmental stages (2-6 months and 7-14 months) during walking. *p *<* 0.05, **p *<* 0.01, ***p *<* 0.001..

### 2.2. Biomechanical data collection

Motion capture, ground reaction force, and subject mass data during walking and trotting gaits were collected. An experienced handler guided subjects at self-selected speeds at the UIUC Large Animal Hospital (Fig. 1B). Subjects were verbally encouraged to increase speed over the course of data collection. Each trial included multiple gait cycles.

A six-camera motion capture system (Qualysis Oqus 500+) was used to record marker positions (sampling frequency of 160 Hz) of 20 anatomical locations on the left side of the foal: 9 markers on the forelimb, 10 on the hindlimb, and one on the head (Fig. 1C). The ground reaction forces of the left forelimb were measured using an in-ground force plate (AMTI Model EQ6001200-400 Inc, Watertown, Massachusetts) at a sampling frequency of 350 Hz.

### 2.3. Data processing

To account for differences in subject size due to normal biological variability and age, we calculated the dimensionless Froude number [Alexander and Jayes, 1983; Alexander, 1996; Hof, 1996] to compare animals moving with dynamic similarity. For each trial, the Froude number (*Fr*) was calculated as

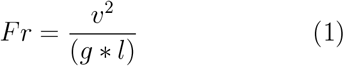

where *v* = absolute velocity (m/s), *g* = 9.81*m/s*^2^, and *l* = limb length (m).

The absolute velocity was calculated using the difference in the shoulder marker position (Fig. 1C) in the direction of movement between the beginning of the stance phase of one cycle and the beginning of the stance phase two strides later, divided by the time of movement (MATLAB, vR2021a). A minimum of three complete strides were used from each trial to calculate the velocity. Limb length was calculated using a single frame during the stance phase in the slowest walk trial in which the joint angles were maximally extended (akin to a natural standing pose) (MATLAB, vR2021a). The difference in the vertical position of the elbow and hoof markers (Fig. 1C) was used to obtain limb length. For each foal, five neutral pose events were averaged to calculate age-specific limb length.

Ground reaction force (GRF) data were filtered using a 2^*nd*^ order low-pass Butterworth filter with a cutoff frequency of 10 Hz. The measured peak forces in the vertical and anterior-posterior (A-P) directions were extracted from each trial. Trials were considered valid when only the left hoof was in full contact with the force plate and the peak GRF fell within the range expected for horses (4-16 N/kg) [Merkens et al., 1986, 1993a; Griffin et al., 2004; Weishaupt et al., 2010; Amitrano et al., 2016; Gorissen et al., 2017].

The Froude numbers of all trials were plotted as a function of age in months to evaluate the gait transition from walking to trotting at relative speeds during growth (R, v4.2.2).

### 2.4. Analytical predictions of GRF

Changes in Froude number with respect to age, velocity, and gait were characterized. We then evaluated two analytical models for predicting GRF data based on the Froude number (Fr). First, a non-linear logistic regression [Valette et al., 2008] fit to all data for each GRF component was assessed:

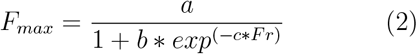

where *F*_*max*_ is peak force, *Fr* is the Froude number, and *a, b, c* are parameters fit to the model.

We also tested a linear regression to individual gait patterns (walk vs. trot) for each GRF component:

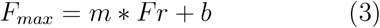

Goodness of fit was characterized using the average absolute percent error (herein referred to as error) and the R-squared value. Statistical significance was based on an alpha = 0.05. To evaluate whether the overall fit is appropriate for different developmental stages, foals were grouped based on growth rate phases (2-6 months, 7-14 months) [Rogers et al., 2021] (R, v4.2.2).

## 3. Results

The number of trials used for analysis was 16 for walking and 148 for trotting.

### 3.1. Walk-trot transition

The Froude number (Fr) ranged from 0.18 to 0.37 during walking and from 0.69 to 3.68 during trotting. The walk-trot transition occurred near Fr = 0.5 (velocity range of 1.75-2.15 m/s) independent of age (Fig. 2A, B).

**Figure 2.**
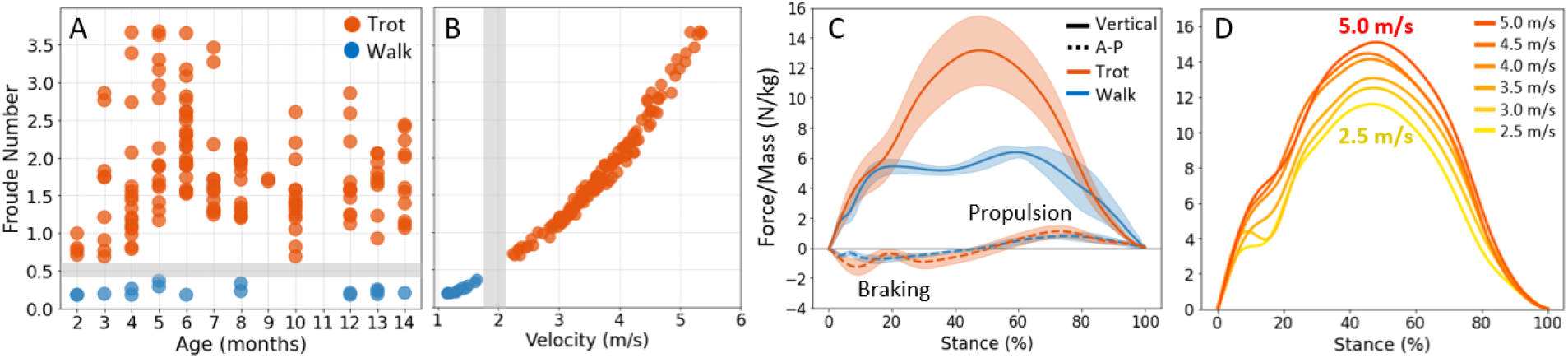
(A) Walk and trot Froude numbers of foals from 2-14 months of age. (B) Froude number as a function of absolute velocity with walk-trot transition illustrated as gray gap. (C) Mean vertical (solid lines) and anterior posterior (dashed lines) ground reaction forces of foals aged 2-14 months (data normalized to body weight). (D) Vertical GRF patterns at different velocities.

### 3.2. Ground reaction forces during walking and trotting

The shape of the vertical ground reaction force (GRF) during walking was bimodal with two distinct peaks corresponding to the braking and accelerating phases, respectively (solid blue line, Fig. 2C). The initial peak occurred at 26.04 ± 7.69% of the stance phase and the second peak occurred at 60.30 ± 9.89% of the stance. There was a moderate increase in the vertical force between the initial peak (5.64 ± 0.47 N/kg) and the second peak (6.61 ± 0.36 N/kg) (Fig. 2C). In contrast, the GRF during trotting had a single peak that occurred at 48.30 ± 2.89% of stance (solid orange line, Fig. 2C). During trotting, the maximum braking and propulsion forces were out of phase with the vertical GRF peak, with the transition from braking to propulsion occurring at 51.84 ± 5.09% stance when the vertical GRF was at its maximum (dashed orange line, Fig. 2C).

The anterior-posterior ground reaction force (A-P GRF) during both walking and trotting started with a negative peak that transitioned to a positive peak around 50 % of the stance phase (48.83 ± 3.80% of stance phase during walking and 51.84 ± 5.09% during trotting), representing the braking and propulsion phases, respectively (Fig. 2C). During walking, the maximum braking forces occurred at 16.57 ± 7.73% and the maximum propulsion forces occurred at 75.74 ± 6.39% of the stance phase (dashed blue line, Fig. 2C). The maximum force was higher during propulsion (0.95 ± 0.31 N/kg) than braking (−0.83 ± 0.12 N/kg) (Table 1). During trotting, the maximum braking forces occurred at 14.10 ± 10.80% and the maximum propulsion forces occurred at 72.97 ± 5.41% of the stance phase, close to walking (dashed orange line, Fig. 2C). However, braking forces were higher (−1.48 ± 0.35 N/kg) than propulsion (1.11 ± 0.29 N/kg) during trotting (Table 1).

**Table 1:** Summary of gait metrics during walking and trotting. R-squared and average absolute percent error (error) of linear and logistic regressions of maximum ground reaction force. *p *<* 0.05, **p *<* 0.01, ***p *<* 0.001.

|  | Velocity (m/s) | Froude | Maximum ground reaction force (N/kg) |  |  |
| --- | --- | --- | --- | --- | --- |
|  |  |  | Vertical | Braking | Propulsion |
| Walk | $1.34 \pm 0.16$ | $0.23 \pm 0.06$ | $6.61 \pm 0.36$ | $-0.83 \pm 0.12$ | $0.95 \pm 0.21$ |
| Trot | $3.71 \pm 0.7$ | $1.81 \pm 0.69$ | $13.16 \pm 1.48$ | $-1.48 \pm 0.35$ | $1.11 \pm 0.29$ |
| Regression models |  |  | Vertical | Braking | Propulsion |
| Walk - Linear |  | R-squared | 0.58*** | -0.05 | 0.28 |
|  |  | Error | 2.72% | 11.94% | 16.63% |
| Trot - Linear |  | R-squared | 0.23*** | 0.28*** | 0.49*** |
|  |  | Error | 7.92% | 17.30% | 14.27% |
| Walk,Trot - Logistic |  | R-squared | 0.68*** |  |  |
|  |  | Error | 8.38% |  |  |

**Table 1:** Summary of mean values with respective standard deviation (s.d.) shown in parenthesis for Froude number, velocity, maximum vertical, maximum propulsion, and maximum braking forces according to age in months and overall.

| Months | 2 |  | 3 |  | 4 |  | 5 |  | 6 |  |
| --- | --- | --- | --- | --- | --- | --- | --- | --- | --- | --- |
| *Trot* | *n=4* |  | *n=9* |  | *n=17* |  | *n=16* |  | *n=26* |  |
| Froude number | 0.82 | (0.12) | 1.61 | (0.81) | 1.62 | (0.86) | 2.26 | (0.86) | 2.26 | (0.55) |
| Velocity (m/s) | 2.40 | (0.18) | 3.34 | (0.85) | 3.39 | (0.82) | 4.08 | (0.76) | 4.16 | (0.51) |
| Max vertical (N/kg) | 12.65 | (1.10) | 14.75 | (0.78) | 12.88 | (1.28) | 14.10 | (1.77) | 13.74 | (1.12) |
| Max propulsion (N/kg) | 0.91 | (0.27) | 1.24 | (0.34) | 1.09 | (0.34) | 1.37 | (0.32) | 1.27 | (0.23) |
| Max braking (N/kg) | -0.99 | (0.41) | -1.43 | (0.34) | -1.36 | (0.28) | -1.76 | (0.42) | -1.67 | (0.35) |
| *Walk* | *n=2* |  | *n=1* |  | *n=2* |  | *n=2* |  | *n=1* |  |
| Froude number | 0.18 | (0.00) | 0.19 | - | 0.22 | (0.06) | 0.33 | (0.05) | 0.18 | - |
| Velocity (m/s) | 1.16 | (0.01) | 1.18 | - | 1.29 | (0.16) | 1.58 | (0.10) | 1.19 | - |
| Max vertical (N/kg) | 6.25 | (0.23) | 6.53 | - | 6.60 | (0.18) | 7.22 | (0.67) | 6.66 | - |
| Max propulsion (N/kg) | 0.89 | (0.33) | 0.73 | - | 0.97 | (0.29) | 1.16 | (0.39) | 0.77 | - |
| Max braking (N/kg) | -0.76 | (0.18) | -1.01 | - | -0.93 | (0.11) | -0.74 | (0.16) | -0.83 | - |
| Months | 7 |  | 8 |  | 9 |  | 10 |  | 12 |  |
| *Trot* | *n=13* |  | *n=15* |  | *n=2* |  | *n=15* |  | *n=11* |  |
| Froude number | 1.82 | (0.73) | 1.70 | (0.37) | 1.70 | (0.02) | 1.46 | (0.50) | 1.74 | (0.57) |
| Velocity (m/s) | 3.74 | (0.68) | 3.65 | (0.41) | 3.70 | (0.03) | 3.38 | (0.57) | 3.75 | (0.59) |
| Max vertical (N/kg) | 14.27 | (0.86) | 11.98 | (0.48) | 13.63 | (0.40) | 11.34 | (1.18) | 13.27 | (1.47) |
| Max propulsion (N/kg) | 1.04 | (0.29) | 1.08 | (0.16) | 1.01 | (0.31) | 0.89 | (0.15) | 1.04 | (0.25) |
| Max braking (N/kg) | -1.62 | (0.23) | -1.41 | (0.21) | -1.75 | (0.17) | -1.17 | (0.27) | -1.44 | (0.26) |
| *Walk* | *n=0* |  | *n=2* |  | *n=0* |  | *n=0* |  | *n=2* |  |
| Froude number |  |  | 0.28 | (0.07) |  |  |  |  | 0.19 | (0.02) |
| Velocity (m/s) |  |  | 1.48 | (0.19) |  |  |  |  | 1.27 | (0.06) |
| Max vertical (N/kg) |  |  | 6.65 | (0.18) |  |  |  |  | 6.43 | (0.24) |
| Max propulsion (N/kg) |  |  | 1.09 | (0.12) |  |  |  |  | 0.94 | (0.12) |
| Max braking (N/kg) |  |  | -0.85 | (0.13) |  |  |  |  | -0.81 | (0.15) |
| Months | 13 |  | 14 |  | Overall |  | % of Max vertical |  |  |  |
| *Trot* | *n=9* |  | *n=11* |  | *n=148* |  |  |  |  |  |
| Froude number | 1.67 | (0.38) | 1.73 | (0.52) | 1.81 | (0.69) | - |  |  |  |
| Velocity (m/s) | 3.80 | (0.46) | 3.76 | (0.58) | 3.71 | (0.70) | - |  |  |  |
| Max vertical (N/kg) | 12.15 | (0.54) | 13.17 | (0.96) | 13.16 | (1.48) | - |  |  |  |
| Max propulsion (N/kg) | 0.98 | (0.11) | 0.98 | (0.19) | 1.11 | (0.29) | 8.43 |  |  |  |
| Max braking (N/kg) | -1.30 | (0.21) | -1.56 | (0.26) | -1.48 | (0.35) | 11.02 |  |  |  |
| *Walk* | *n=3* |  | *n=1* |  | *n=16* |  |  |  |  |  |
| Froude number | 0.22 | (0.03) | 0.21 | - | 0.23 | (0.06) | - |  |  |  |
| Velocity (m/s) | 1.40 | (0.09) | 1.31 | - | 1.34 | (0.16) | - |  |  |  |
| Max vertical (N/kg) | 6.52 | (0.32) | 6.80 | - | 6.61 | (0.36) | - |  |  |  |
| Max propulsion (N/kg) | 0.95 | (0.12) | 0.72 | - | 0.95 | (0.21) | 14.37 |  |  |  |
| Max braking (N/kg) | -0.83 | (0.09) | -0.76 | - | -0.83 | (0.12) | 12.41 |  |  |  |

As expected, the magnitude of the maximum vertical GRF increased with increasing velocity and the overall shape of the curve was preserved (Fig. 2D). The average maximum vertical GRF during trotting (13.16 ± 1.48 N/kg) was nearly twice that of walking (6.61 ± 0.36 N/kg) (Table 1). The maximum braking forces were 1.78-fold higher during trotting than walking whereas the maximum propulsive forces were more similar during trotting and walking (1.16-fold increase).

### 3.3. Analytical predictions of maximum GRFs

The linear regression-based prediction of vertical GRF from Froude number was more accurate (error for walk: 2.72%; trot: 7.92%) than the logistic expression (8.38%) (Table 1, Fig. 3A). A logistic regression could not be used for the maximum propulsive or braking forces because of the discontinuity in forces between the walk and trot gaits (Fig. 3B,C).

**Figure 3.**
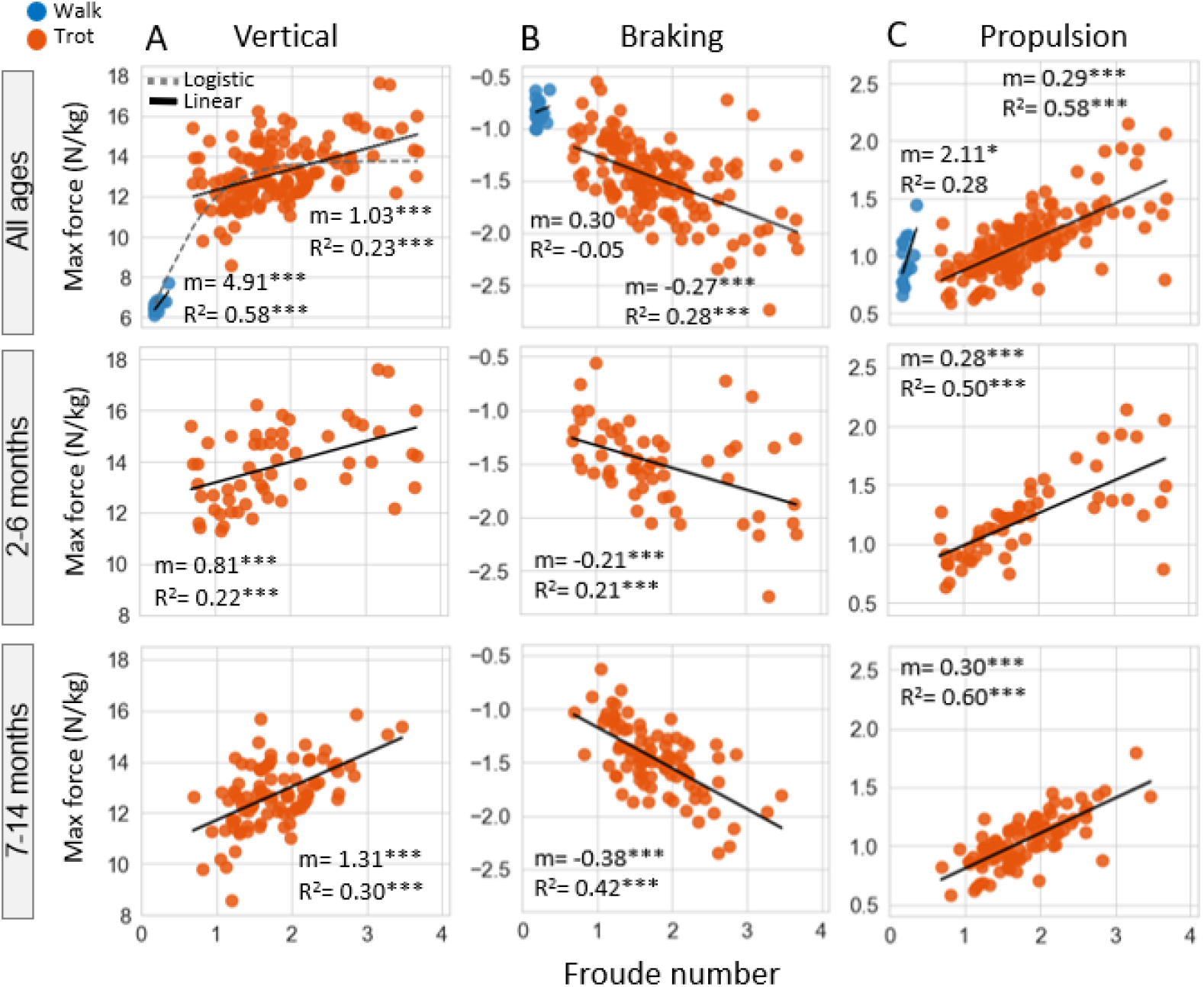
(A) Linear and non-linear logistic models compared to analyze the relationship between maximum vertical GRF (N/kg) and Froude number with and without gait distinction, respectively. Maximum propulsion (B) and braking (C) (max and min A-P GRF, respectively) relation to Froude Number analyzed using linear fit by gait. Logistic regression equation for maximum vertical forces: = 13.77*[1 + 0.21\**exp*(*−*0.41 * *Fr*)]^*−*^1.

Predictions of walking ground reaction force were most accurate for the vertical GRF (error = 2.72%), followed by braking (error = 11.94%), and then propulsive GRF (error = 16.63%) (Table 1). Errors in ground reaction force predictions during trotting were higher than those during walking. The maximum vertical GRF during trot had the lowest average absolute error (7.92%), followed by propulsive forces (14.27%) and then braking (17.30%) (Table 1). The amount of variability in the maximum ground reaction force that could be explained by Froude number was greatest for the vertical GRF during walking (R^2^ = 0.58, p*<*0.001) while the correlation between propulsive or braking forces and Froude number was not significant. All linear correlations between Froude number and the ground reaction forces during trotting were significant (p*<*0.001). Based on the linear regressions, the maximum vertical forces increased at a higher rate with increasing Froude number during walking than trotting (Fig. 3A). During trotting, the maximum propulsion forces increased at a slightly higher rate with increasing Froude number than the increase in braking forces (Fig 3B, C).

Finally, we compared the age-specific relationship between Froude number and ground reaction forces during trotting. Overall, the ground reaction force was more variable between two and six months of age with the greatest variability occurring in the vertical component (Fig. 3A). The relationship to Froude number and GRFs was consistently stronger in the 7-14 month age group, in particular for the braking and propulsive phases. The rate of increase in GRF with increasing Froude number was higher for the 7-14 month group compared to the 2-6 month groups (Fig. 3B,C).

## 4. Discussion

Although direct measurement of ground reaction forces (GRFs) using force plates, instrumented treadmills or pressure sensitive insoles would be preferred, there remain challenges in either accessibility or application in natural ground conditions, which limit field-based studies of locomotion. Motivated by the intrinsic coupling between kinematics and kinetics dictated by the dynamic equations of motion, we explored a simple analytical model for predicting the GRFs during equine gait. Importantly, for ontogenetic studies, accounting for variation in size, age, and speed during gait is critical as normalizing GRFs by mass alone is not sufficient [Voss et al., 2010]. We therefore based our model on the Froude number to allow for comparing animals of different sizes [Alexander and Jayes, 1983; Voss et al., 2010; Mölsä et al., 2010; Abdelhadi et al., 2013] including horses [Griffin et al., 2004; Weishaupt et al., 2010; Gorissen et al., 2017].

Foals in this study were locomoting at self-selected speeds. Foals walked at 1.34 ± 0.16 m/s (Fr=0.23 ± 0.06) and trotted at 3.71 ± 0.7 m/s (Fr=1.81 ± 0.69). Hoyt et al [Hoyt and Taylor, 1981] showed that self-selected speeds within each gait occurred at energetically optimal values (1-1.50 m/s for walking and 3-3.75 m/s for trotting). We found that foals transitioned from a walk to a trot at Froude numbers lower than 0.5 (Fig 2A) and was consistent for all ages studied (2-14 months). In adult horses, the walk-trot transition has been reported to occur at 0.35 ± 0.03 [Griffin et al., 2004] which is similar to the highest Froude number we found in growing horses (Fr = 0.37). It is widely known that animals and humans transition between gaits to minimize the total metabolic cost of transport [Hoyt and Taylor, 1981; Cavagna et al., 1977]. Our data suggests that optimizing the cost of transport occurs early in life. Unlike other mammals, horses begin standing and walking within hours of birth and it may be that their musculoskeletal system is genetically primed to transition from walking-trotting without requiring an adaptation period to locomotion.

With respect to the ground reaction force profiles in growing horses, our data mimic the reports of GRF profiles in adult horses [Merkens et al., 1986, 1993a,b; McLaughlin et al., 1996; Dutto et al., 2004; Weishaupt et al., 2010]. To our knowledge, only two studies have reported GRFs in young horses [Amitrano et al., 2016; Gorissen et al., 2017]. Amitrano et al. [Amitrano et al., 2016] measured the maximum vertical, braking, and propulsion GRFs in six Standardbred horses of ages 11-14 months during walking (6.74 ± 0.66 N/kg, −0.95 ± 0.18 N/kg, 0.95 ± 0.20 N/kg, respectively). These values are similar to our measurements of the maximum vertical, braking, and propulsion forces across ages 12-14 months during walking (6.54 ± 0.27 N/kg, −0.81 ± 0.01 N/kg, 0.91 ± 0.13 N/kg, respectively). Gorissen et al. [Gorissen et al., 2017] conducted a longitudinal study of Royal Dutch Sport Horse foals (n=11) during the first 6 months of life. During trotting (Fr range: 0.55-0.78), the average maximum vertical force was 14.04 ± 1.20 N/kg which is similar to our results of 13.80 ± 0.93 N/kg during trotting (Fr range: 0.69-0.77). The close similarity in these maximum vertical force values suggests consistency in normalized ground reaction force (GRF) measurements across breeds, supporting the reliability of GRF predictions in different horses.

The maximum propulsion forces during walking were higher than the maximum braking forces, whereas during trotting the maximum braking forces were higher than maximum propulsion forces (Table 1). The increased speeds of trotting would intuitively suggest the need for greater braking force (1.78-fold higher than walking). Interestingly, at trotting speeds (3.71 ± 0.7 m/s) horses needed just 1.16-fold higher propulsion forces compared to walking suggesting a conservation of energy that allows for increasing speed with less propulsive forces. Energy storage may be facilitated by the presence of the long tendons of the lower limbs which can return energy to the system following initial stretch [Biewener, 1998].

The rate of change of ground reaction forces with increasing Froude number was activity dependent. Maximum vertical and propulsive GRFs increased at a higher rate during walking than trotting. However, the range of Froude numbers during walking is small (0.23 *±* 0.06) and the range of maximum forces during walking was similarly small. It is debatable whether the linear regressions of normalized force with respect to Froude number are in fact physiologically relevant. In contrast, the increasing rate of GRF during trotting was more appreciable. Vertical GRF increased by 1 N/kg per unit increase in Froude number. The rate of increase in the braking and propulsive forces was approximately 1/3 that of the vertical GRF. The increase in rate of vertical GRF with increasing speed is likely due to increased flexion of the lower limbs that contribute to increased downward acceleration of the center of mass during trotting.

The prediction of maximum ground reaction force values for a given Froude number was more accurate for linear regression models compared to a logistic regression model. The ground reaction forces in foals younger than six months of age were more variable than the GRFs in foals older than seven months. This variability was more pronounced when the Froude number was greater than three. The variability in young foals could be reflective of the time it takes to learn how to move consistently at higher speeds. Nonetheless, the largest errors occurred when predicting the braking and propulsion maximum magnitudes. These errors could be attributed to individual variation in gait, which are not captured by the Froude number. Factors such as phase-specific kinematics and dynamics, including differences in limb velocity profiles during the stance and swing phases, variations in foot strike patterns and limb posture at contact, and changes in body mass distribution and limb proportions during growth, all contribute to the variability in these forces.

There are some limitations to this study. The self-selected speeds resulted in a limited range of speeds that could be studied and a low number of walk trials available for analysis. The range of speeds presented is a result of the foals’ natural gait within the confines of a large animal veterinary hospital hallway. The sample size in our study is relatively small and data from 0-2 months of age are missing. Lastly, although the model is useful for GRF predictions over a range of Fr numbers, errors in braking and propulsion were 12% on average. However, the largest contribution to the ground reaction force is the vertical GRF for which our analytical models had errors less than 3%.

While the analytical predictions are not error free, they do provide an estimate of the ground reaction forces during walking and trotting gaits which could be useful for studies of cohorts of horses locomoting at different speeds or of different ages. We used the linear regression models to create a Python package (v3.10) that can output subject-specific ground reaction force profiles based on the subject velocity and limb length. Specifically, a representative curve of the vertical and anterior-posterior ground reaction force was normalized to the maximum GRF value serving as a GRF template. Following calculation of the Froude number, the peak values of the ground reaction force predicted by the analytical model during trotting and walking for each degree of freedom were then used to scale the ground reaction force template (Fig. 4). Froude numbers below 0.50 are considered a walking gait, while values greater than 0.50 are considered a trotting gait.

**Figure 4.**
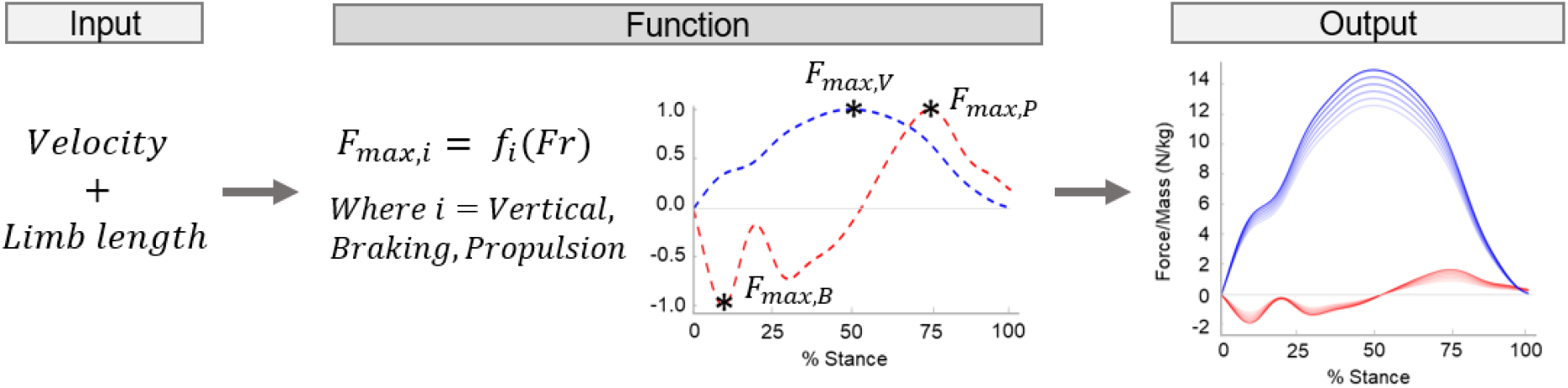
Pipeline summary of code that creates a subject- and trial-specific set of ground reaction forces. A trot trial is shown as an example with varying absolute velocities.

To our knowledge, this is the first study to develop a predictive analytical model of ground reaction forces in foals that includes a comprehensive assessment of foals younger than 12 months of age. Our development of an analytical model for predicting GRFs based on limb length and velocity now enables GRF predictions in real world settings. These data can be paired with in-field markerless motion capture or treadmill exercise to estimate internal joint moments and muscle forces. The capacity to collect such data will enable further studies of natural locomotion in horses with implications for studies of exercise and other musculoskeletal diseases.

## 5. Acknowledgments

The authors are grateful to Griffin Sipes and Dr. Kellie Halloran who assisted with data collection and code development.

## 7. Supplemental Data

**Linear - Walk:**

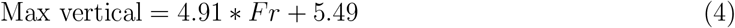

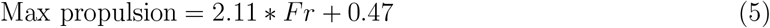

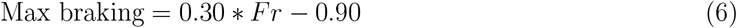

**Linear - Trot:**

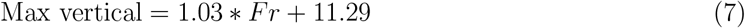

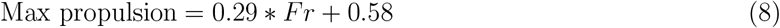

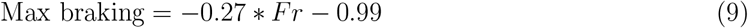

